# High Fat Diet Stimulates Beta-Oxidation, Alters Electrical Properties and Induces Adipogenicity of Atria in Obese Mice

**DOI:** 10.1101/2020.10.20.347161

**Authors:** Nadine Suffee, Elodie Baptista, Jérôme Piquereau, Maharajah Ponnaiah, Nicolas Doisne, Farid Ichou, Marie Lhomme, Camille Pichard, Vincent Galand, Nathalie Mougenot, Gilles Dilanian, Laurence Lucats, Elise Balse, Mathias Mericskay, Wilfried Le Goff, Stéphane Hatem

## Abstract

Metabolic disorders such as obesity are risk factors of atrial fibrillation, not only by sharing comorbidities but likely through their direct impact on atria, notably its adipogenicity. Here, we submitted mice that lack cardiac adipose tissue to a high fat diet and first studied the atrial metabolomic and lipidomic phenotypes using liquid chromatography-mass spectrometry. We found an increased consumption of free fatty acid by the beta-oxidation and an accumulation of long-chain lipids in atria of obese mice. Free fatty acid was the main substrate of mitochondrial respiration studied in the saponin-permeabilized atrial muscle. Conducted action potential recorded in atrial trabeculae was short, and ATP-sensitive potassium current was increased in perforated patch-clamp atrial myocytes of obese mice. There was histological and phenotypical evidence for an accumulation of adipose tissue in obese mice atria. Thus, an obesogenic diet transforms the energy metabolism, causes fat accumulation and induces electrical remodeling of atria myocardium.

**HIGHLIGHTS:**

- Untargeted metabolomic and lipidomic analysis revealed that a high fat diet induces profound transformation of atrial energy metabolism with beta-oxidation activation and long-chain lipid accumulation.
- Mitochondria respiration studied in atrial myocardial trabecula preferentially used Palmitoyl-CoA as energy substrate in obese mice.
- Atria of obese mice become vulnerable to atrial fibrillation and show short action potential due to the activation of K-ATP dependent potassium current.
- Adipocytes and fat molecular markers were detected in atria of obese mice together with an inflammatory profile consistence with a myocardial accumulation of fat.

## INTRODUCTION

Metabolic disorders such as obesity, metabolic syndrome or diabetes are risk factors for atrial fibrillation (AF), the most frequent cardiac arrhythmia in clinical practice (Dublin et al., 2010; Huxley et al., 2012; Johansen et al., 2008). For instance, each increase in body mass index (BMI) is associated with 4% increase in incidence of AF. Indeed, obesity is now considered an arrhythmogenic risk factor (Johansen et al., 2008). There is also a correlation between the duration of diabetes mellitus, the level of dysglycemia and the risk of AF (Bohne et al., 2019).

Whether this epidemiologic association between AF and metabolic disorders is due only to shared comorbidities or also to causal pathophysiological links remains to be established. In this vein, a number of articles report an impact of altered myocardial metabolism on the development of AF. For instance, transcriptomics and metabolomics analysis of human atrial samples shows that AF causes a major metabolic stress of the myocardium with the predominant use of glucose and ketone bodies as the main feature (Chao et al., 2010; Lai et al., 2018; Latini et al., 2013). An upregulation at both transcript (Barth et al., 2005) and protein (Mayr et al., 2008) levels of several glycolytic enzymes has been observed in atrial appendage from patients with AF, suggesting that glycolysis overcomes the short-term energy demand in AF atrium. Moreover, in dilated and fibrotic atria associated with heart failure, pyruvate dehydrogenase and glucose oxidation are reduced (De Souza et al., 2010). A relationship between dysmetabolism and arrhythmia is also supported by the vulnerability to AF observed in several experimental models of obesity and metabolic syndrome (Iwasaki et al., 2012).

A potential interface between AF and metabolic disorders is the epicardial adipose tissue (EAT), an abundance of which is associated with both increased incidence of AF and poor outcome after an ablation procedure of the arrhythmia (Al Chekakie et al., 2010; Thanassoulis et al., 2010; Wong et al., 2011). EAT is an important source of free fatty acid (FA) and of adipokines; it regulates the redox state of the atrial myocardium (Antonopoulos et al., 2016; Villasante Fricke and Iacobellis, 2019). However, under certain clinical circumstances, EAT can secrete pro-fibrotic factors and cytokines that contribute to the arrhythmogenic fibro-fatty remodeling of the atrial myocardium (Haemers et al., 2017; Nalliah et al., 2020). The mechanisms regulating the accumulation of EAT in the atria are still poorly understood. Obese patients show an accumulation of EAT at the surface of the atria suggesting a link between metabolic disorders and myocardial adipogenicity (Mahajan et al., 2018).

Here we tested the hypothesis that, through modifications of metabolomics and lipidomic phenotypes of the myocardium, an obesogenic diet can be an adipogenic factor for atria. Mice that lack cardiac adipose tissue were submitted to a prolonged high fat diet, and the atrial myocardium was studied using lipidomic and metabolomic approaches, mitochondria respiration assay and cellular electrophysiological measurements. We found that a high fat diet causes a profound transformation of the functional and histological properties of the atrial myocardium that become fat and vulnerable to AF.

## METHODS

### Study approval

All animal experiments were in compliance with the *Guide for the Care and Use of Laboratory Animals*, according to the Directive 2010/63/EU of the European Parliament approved by the local committee of animal care (Agreement A751320).

### Experimental model and clinical study

Eight-week-old male C57Bl/6J mice (20-25g) purchased from Janvier (CERJ, Laval, France) were maintained under a 12-hour light/12-hour dark cycle at a constant temperature (21°C) with free access to food and water. The mice were subjected to 4 months of either a high fat diet (HFD, 60% fat, D12492i Research diet) (n=60) or a normal diet (ND, 8,4% fat, A04 Safe) (n=60).

### Clinical study

In mice under 0.2-0.5 % isoflurane anesthesia, transthoracic echocardiography was performed for cardiac examination using a Vivid 7 dimension cardiovascular ultrasound system equipped with a probe of 9-14 MHz frequency (GE Healthcare, Vélizy, France) as described <u>in the supplemental methods herein</u>. Also in anaesthetized mice, atrial burst pacing was delivered through an esophageal probe (see the <u>supplemental methods</u>).

### Insulin Test Tolerance (ITT), Glucose Tolerance Test (GTT)

At 16-weeks diet endpoint, mice were fasted for 6 hours before GTT or ITT. The GTT was performed using a 2 g/kg glucose bolus (Sigma-Aldrich) injected in intraperitoneal cavity (n=10, each group). The ITT was performed using insulin (0.75 U/kg body weight) (Novo Nordisk) injected intraperitoneally (n=10, each group). For both tests, blood samples were collected from tail vein at 0, 15, 30, 60, 90, and 120 minutes post-injection, and glucose concentrations were determined using a glucose meter (Accu-Chek Performa, Roche).

### Insulin assay

At 16 weeks of HFD or ND, blood mice samples were collected and immediately centrifuged. The plasma was separated and stored at −80°C for insulin assay according to manufacturer’s instructions (Roche).

### Electrophysiological measurements

#### Action Potential Recording

Left atria were excised rapidly from mice and continuously superfused in Tyrode’s solution containing (mM):NaCl 120; KCl 5.6; NaH_2_PO_4_, H_2_O 0.6; NaHCO_3_ 24; MgCl_2_, 6H_2_0 1.1; CaCl_2_, 2H_2_0 1.8 supplemented with glucose 2g/L and sodium pyruvate 200mg/L and equilibrated with 95% O_2_/5% CO_2_ at 37 ± 0.5°C (pH 7.4). Tissues were stimulated (Isostim A320, World Precision Instrument, Inc.) with a rectangular pulse of 2-ms at twice threshold through a bipolar Ag-electrode and were allowed to equilibrate for at least 1 hour at 1Hz. Action potentials (APs) were recorded using glass microelectrodes, filled with 3 M KCl and having tip resistances over 20 MΩ, coupled to high-input impedance preamplifier (Intracellular Electrometer IE-210, Warner Instrument Corp). APs were displayed on an oscilloscope and simultaneously digitized through Digidata 1200 (Axon Instruments) and analyzed with Acquis1 software, version 4.0 (Gerard Sadoc, CNRS, Gif/Yvette, France). AP parameters included action potential duration (APD) at 75 and 90% of repolarization.

The pharmacological study on K-ATP modulators on action potential was performed by superfusing atrial trabeculae with Tyrode’s solution supplemented with 30 μM cromakalim (Sigma-Aldrich) or 1 μM glibenclamide (Sigma-Aldrich). APs were recorded before and after drug application.

#### Potassium current recording

Ionic currents were recorded in enzymatically isolated atrial myocytes as previously described (Boycott et al., 2013). Small pieces of tissue were briefly dissociated in Tyrode’s solution with low calcium, low magnesium (in mM: NaCl 140; KCl 5.4; MgCl_2_ 0.5; CaCl_2_ 0.07; HEPES 5; glucose 5.5; KH2PO_4_ 1.2; taurine 50; pH 7,4) supplemented with 3mg/mL BDM, 1mg/mL bovin serum albumin (BSA), collagenase, elastase and protease at 37°C. Isolated myocytes were then collected in KB solution containing (mM) (KCl 25; MgCl_2_ 3; HEPES 10; glucose 10; KH_2_PO_4_ 10; taurine 20; EGTA 0.5; L-glutamic acid 70; pH 7.4) warmed at 37°C. Currents were recorded using an Axopatch 200B amplifier (Axon Instruments) and patch pipettes (Corning 7052 Kovar Sealing, World Precision Instruments) with a resistance between 1 and 2MΩ. The extracellular Tyrode’s solution used for current recording contained (mM): NaCl 135; KCL 4; MgCl_2_ 2; CaCl_2_ 2; HEPES 10; glucose 20; Na_2_HPO_4_ 1, sodium pyruvate 2.5 (pH 7.4) supplemented with 1 μM ryanodine. The intracellular pipette solution used for whole cell recording contained (mM): NaCl 10; KCl 10; MgCl_2_ 3; HEPES 10; glucose 10; K-aspartate 115, and EGTA 10 (pH 7.2) for the perforated patch-clamp (mM): KCl 20; MgCl_2_ 1; HEPES 10; K-aspartate 110 (pH 7.2) and 250μg/mL nystatin for perforated patch-clamp. For the current-voltage membrane potential (V_m_) and activation-V_m_ protocol, currents were elicited by 500-ms depolarizing pulses at 0.2 Hz from a holding potential of −80 mV, in 10 mV increments, between −130 and +60 mV. To study K_ATP_ and K_1_ currents, 1 μM of glibenclamide (Sigma-Aldrich) and 500 μM of barium were respectively added in perfusion medium. I_K,ATP_ was considered as glibenclamide-sensitive current and IK1 as barium-sensitive current.

### Metabolomic and lipidomic analysis

Food (normal diet and high fat diet), plasma and pooled atria (right and left) were collected from mice after 4 months HFD (n=50) or Normal Diet (n=50), and lipidomics and metabolomics analysis were processed as described in the supplemental methods and Table S1.

### Mitochondrial respiration

Mitochondrial respiration was studied in situ in saponin-permeabilized atrial muscle fibers using a Clarke electrode (Sanz et al., 2019). Fibers were dissected under a binocular microscope, from right and left atria in skinning (S) solution (2.77 mM CaK2 ethyleneglycol tetraacetic acid (EGTA), 7.23 mM K2EGTA, 6.56 mM MgCl_2_, 5.7 mM Na_2_ adenosine triphosphate (ATP), 15 mM phosphocreatine, 20 mM imidazole, 20 mM taurine, 0.5 mM dithiothreitol (DTT) and 50 mM K-methanesulfonate, pH 7.1 at 4°C and permeabilized in the same solution with saponin (50 μg/mL) for 30 minutes at 4°C. Permeabilized fibers were first washed in (S) solution for 10 minutes at 4°C. The permeabilized fibers were then incubated in respiration (R) solution (2.77 mM CaK2EGTA, 7.23 mM K2EGTA, 1.38 mM MgCl_2_, 3 mM K_2_HPO_4_, 20 mM imizadole, 20 mM taurine, 0.5 mM DTT, 90 mM K-methanesulfonate, 10 mM Na-methanesulfonate, pH=7.1) with 2mg/mL of BSA (bovin serum albumin) at room temperature for 10 minutes and transferred into 3 mL oxygraphic chambers (Strathkelvin Instruments, Glasgow, UK). Oxygen consumption was measured after successive addition of ADP (2 mM), malate (4 mM), palmitoyl-CoA and carnitine (100 μM and 2 mM), pyruvate (1 mM), glutamate (10 mM), succinate (15 mM), amytal (1 mM), and N,N,N’,N’-Tetramethyl-p-phenylenediamine dihydrochloride (TMPD)-ascorbate (0.5 mM) to R solution with 2mg/mL of BSA (bovin serum albumin) at 23°C. After the experiments, fibers were collected, dried and weighed. Respiration rates were given in μmoles O_2_/min/g dry weight.

### Enzyme activity

Frozen atrial samples were homogenized with a Bertin Precellys 24 homogenizer (Bertin, Montigny-le-Bretonneux, France) in an ice-cold buffer (50 mg/mL) containing: 5 mM HEPES (pH 8,7), 1 mM EGTA, 0,1% Triton X-100 and 1mM DTT. Rates were given as international units (IU)/g protein.

### Histology and immunofluorescence

Mouse hearts were removed (n=10 per group), perfused through the aorta with PBS, fixed in 4% paraformaldehyde (PFA) overnight and were fixed, dehydrated and paraffin-embedded. Seven μm-thick sections of atria samples were stained with Masson’s trichrome according to manufacturer’s instructions (Sigma-Aldrich). Some sections were incubated with antibodies as described in the supplemental methods.

### Adipogenic transcriptomics profile

Total RNA was isolated from mouse atria tissue and was extracted using TRIzol reagent according to the manufacturer’s instructions (Thermo Fisher Scientific). RNA concentration was measured with a NanoDrop spectrophotometer (Thermo Fisher Scientific), and 1 μg was submitted to reverse transcription using superscript III (First Strand cDNA Synthesis Kit; Thermo Fisher Scientific). qPCR was performed using a TaqMan Gene Expression Assay (Thermo Fisher Scientific) as described previously (Suffee et al., 2017). Briefly, 100 ng of cDNA was used for real-time PCR with specific TaqMan (Table S2) and Master Mix primers. Reactions were performed in single 96-well plates in duplicate using StepOnePlus (ABI, Thermo Fisher Scientific). Expression relative to *Gapdh*, and *18S* was calculated using the 2-(ΔΔCt) method. Levels were expressed in arbitrary units (AU) and were normalized to the normal diet.

### Statistical analysis

Data are expressed as means ± SEM. Differences were investigated using the appropriate Mann Whitney t-test or one-way ANOVA and a Bonferroni post hoc analysis and were considered significant at P < 0.05. Statistical analysis was performed with GraphPad Prism 6.0 (GraphPad Software, Inc.). Statistical analyses of both the metabolomics and lipidomics data were performed using Multi Experiment Viewer (MeV) statistical software package (version 4.9.0; http://www.tm4.org/mev/) (Saeed et al., 2003). Data are shown as mean ± SEM. Comparisons of the two groups (ND and HFD) were performed by a two-tailed student’s t-test. Features were considered significant when the p value was below 0.05 after Benjamini-Hochberg for false discovery rate (FDR) correction (Klipper-Aurbach et al., 1995). The volcano plot depicting the enrichment and depletion of metabolomics and lipidomics features together with the fold change were plotted using EnhancedVolcano R package. We used Microsoft Excel to make the forest plot of individual metabolites and lipids (p value <0.05), the fold-change plots of the lipid species alterations (p value < 0.05) and the mountain plots of lipid and metabolites (p value < 0.05). The corrplot R package was used to build the correlation matrix by Pearson parametric correlations with p value < 0.05 as significant.

## RESULTS

### High fat diet induces obesity, glucose intolerance and atrial vulnerability to arrhythmia

Mice under a prolonged, high fat diet gained weight and became obese at 16-weeks (46.93 ± 0.70 g and 28.74 ± 0.34 g in HFD *vs* ND, respectively, P <0.0001, n=30/group) (Figure 1A). Furthermore, at 16 weeks, HFD mice displayed higher fasting glucose and insulin concentrations and exhibited altered glucose tolerance and insulin sensitivity signaling insulin resistance compared to ND mice (Figure 1B-D).

**Figure 1.**
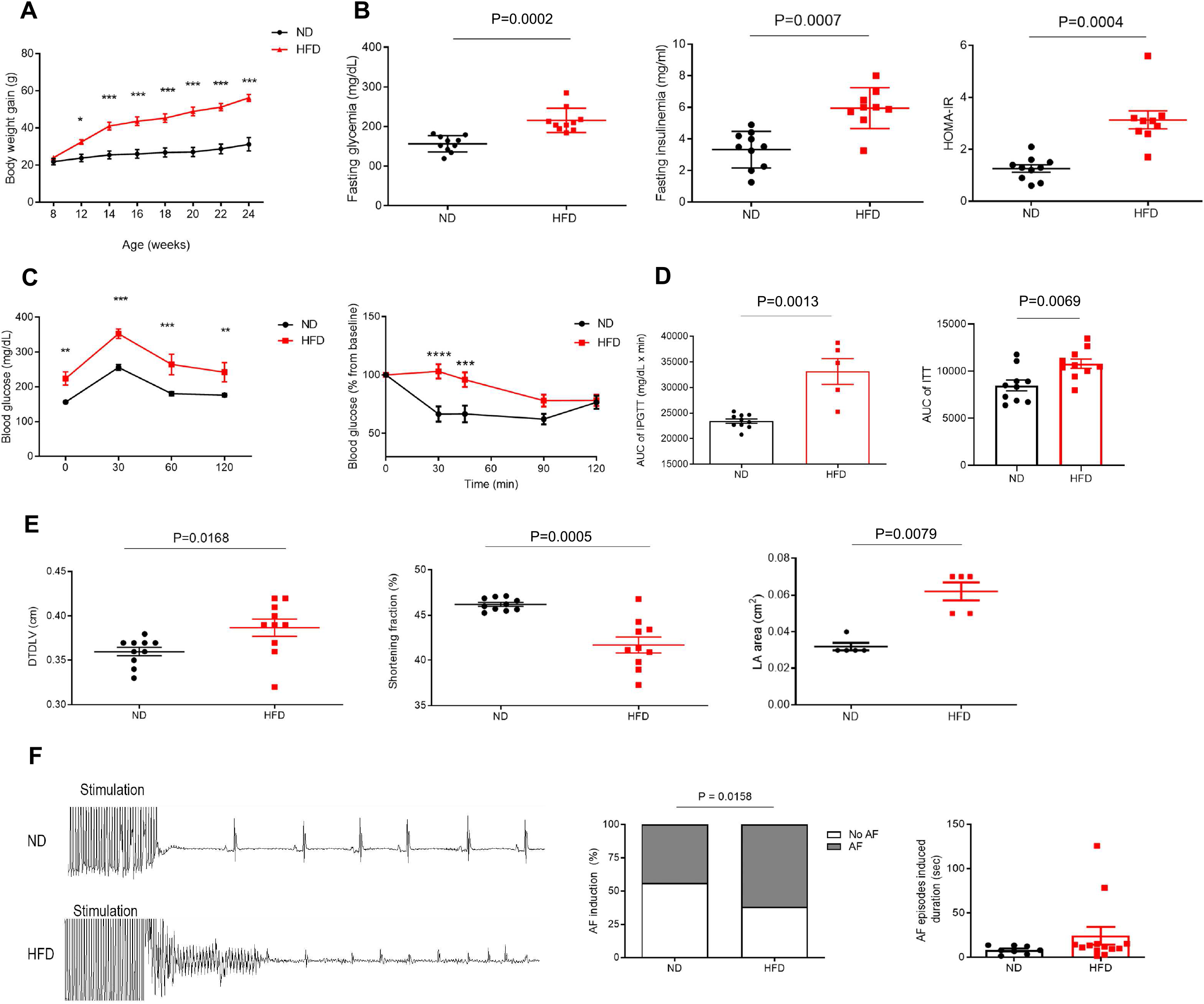
High fat diet induces obesity, glucose intolerance and vulnerability to atrial arrhythmia. (**A**) Body mass of C57BL/6J mice fed with either HFD (n=10) or ND (n=10) for 16 weeks. Data are expressed as mean ± SEM 2-way-ANOVA with Bonferroni’s post-hoc test. *P=0.0476, ***P=0.0002. (**B**) Blood glucose and insulinemia and HOMA-IR index measured at 6h-fasting C57BL/6J mice fed with either HFD (n=10) or ND (n=10) for 16 weeks. (**C**) Time course and (**D**) area under the curve (AUC) of ITT and GTT obtained in HFD (n=5) and ND (n=10). (**E**) Diastolic left ventricle dimeter (DTDLV), shorting fraction and left atria surface (LA) measured using echocardiography in HFD (n=10) or ND (n=10) at 16 weeks. (**F**) Surface electrocardiogram recorded in anesthetized HFD and ND mice following burst of rapid atrial pacing; arrhythmic episodes were quantified as percentage of induction (histogram) and duration (dots). Data are expressed as mean ± S.E.M. Mann–Whitney U test.

Echocardiographic examination performed at 16 weeks showed a moderate dilatation of the left ventricle together with a slight decreased systolic function and atrial dilation in HFD mice (0.032 ± 0.002 cm^2^ *vs* 0.062 ± 0.005, p=0.0079; for HFD or ND mice, respectively n=5/group) (Figure 1E). There was no significant change in heart and atrial weight in obese mice at sacrifice. On electrocardiogram, P wave was slightly prolonged (14.08 ± 1.49 vs 16.04 ± 1.28, p<0.0001), and R-R interval was shorter in 16-week HFD mice. Esophageal delivery of burst of rapid stimulation triggered an episode of AF more frequently and of longer duration in obese than lean, anaesthetized mice (Figure 1F). Taken together, these results indicate that long-lasting HFD induces a metabolic syndrome together with a moderate cardiac remodeling and vulnerability to AF.

### Alteration of the metabolic profile of the atrial myocardium after prolonged high fat diet

Lipidomics and metabolomics profiling of the atrial myocardium as assayed by principal component analysis (PCA) revealed a distinct metabolic profile between groups at 16 weeks (Figure 2A). The volcano plot showed a reduction of carnitine (C0), short-chain acylcarnitines (C2, C3) and short-chain triglycerides (TG), mostly C16, containing fatty acids (48:0, 48:1 and 48:3), whereas glycolysis intermediates (hexose-6-phosphate, hexose-1-phosphate), hexosamine phosphate and long-chain, mostly C18, fatty acid phospholipids (PL) (PC(38:2), PC(40:4), PC(40:5) and PG(36:2), plasmalogens (PE(18:0p/20:4), PE(18:0p/20:5), PE(18:0p/22:6)) and sphingomyelins (SM(36:1), SM(36:2), SM(42:4)) were increased upon HFD (Figure 2B).

**Figure 2.**
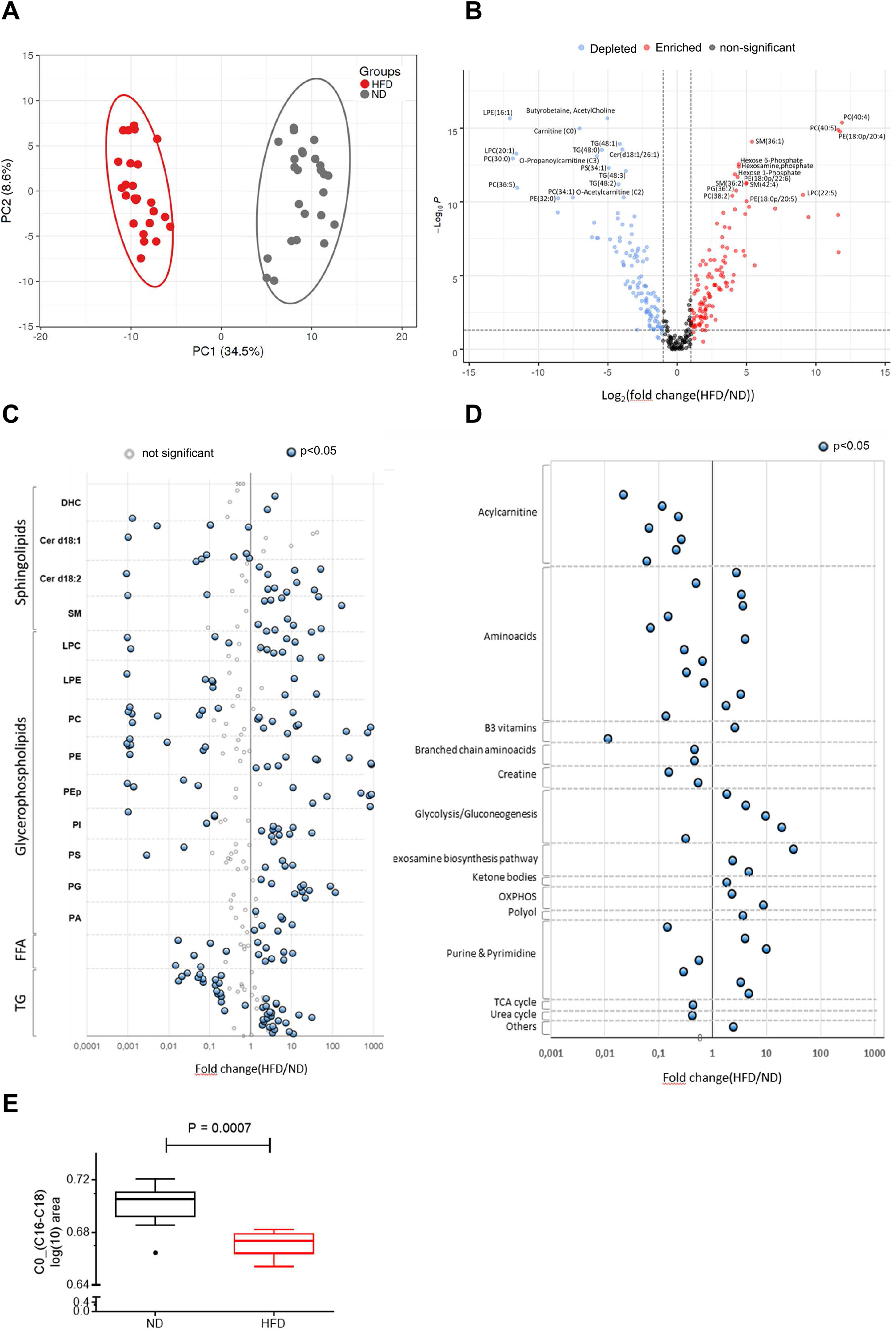
Alteration of the metabolic profile of the atrial myocardium after prolonged high fat diet. (**A**, **B**) Metabolomics and lipidomics features of the atrial myocardium from HFD and ND mice after 16-week of diet (n=25) (**A**) PCA score plot and (**B**) Volcano plot; blue, metabolites depleted; red, metabolites enriched; grey, no significant difference. P < 0.05, fold change ? 1. (**C**, **D**) Forest plots represent individual lipids classified with respect to increasing chain length for each lipid subclass and metabolites (n=25 mice in the two groups). Plots in blue represent individual (**C**) lipids or (**D**) metabolites distinct between HFD and ND conditions. (**E**) Histogram represents (C0) / acylcarnitine (C16) + acylcarnitine C18 ratio measured in mice atrium upon 16-we HFD compared to ND. Data are expressed as mean ± SEM of independent experiments. Groups are compared with t test, P value < 0.05 corrected by FDR, Mann–Whitney test.

Forest plot analysis of distribution of individual lipid species according to lipid classes revealed that 218 lipid species (36 sphingolipids, 122 phospholipids, 18 FFA and 42 TG) were significantly altered in response to the HFD (Figure 2C) and that a majority of them (over 60% of total lipid species) were enriched in atria of HFD mice (Figure 2C). Namely, sphingolipids and the negatively charged PL, PA and PG, showed most of their molecular species enriched in HFD atria independently of the chain length (Supplemental Figure 1A-B). On the contrary, for the other lipid classes (FFA, LPC, LPE, PC, PE, PE plasmalogens, PS, PI and TG) (Supplemental Figure 1C-E), alterations were chain-length dependent with enrichment in long and polyunsaturated and depletion in saturated and short (C16 containing lipids) species. Overall, the atrial myocardium of HFD mice was enriched in C18-containing lipid species as indicated by the reduction of the C16/C18 ratio in all lipid classes and by the increase in total mass of TG and FFA (Supplemental Figure S2A-C).

49 metabolites were modulated by HFD (26 down-regulated and 23 up-regulated) (Figure 2D) including acylcarnitines, branched-chain amino acids (BCAA), creatine and one TCA intermediate were reduced in atrial myocardium while glycolysis and gluconeogenesis intermediates, hexosamine, ketone bodies and polyols were widely increased. Furthermore, at 16 weeks of HFD, transcripts for key enzymes involved in glycolysis and lipogenesis, *i.e*. hexokinase (*Hk2*), acetyl-CoA carboxylase alpha (*Acaca*) and fatty acid synthase (*Fasn*) were increased, whereas diacylglycerol O-acyltransferase homolog 2 (*Dgat2*) was reduced (Supplemental Figure S3A). However, the decrease of the pyruvate/lactate ratio in the atrial myocardium, which mainly reflects the drop of pyruvate, (Supplementary Figure S3B) provided evidence for an impairment of the glycolysis, favoring auxiliary pathways such as hexosamine biosynthesis pathway.

A more detailed analysis of acylcarnitine species brought to light that the carnitine (C0)/acylcarnitine (C16+C18 species) ratio, which is inversely associated with the palmitoyltransferase-1 (CPT-1) activity (Knottnerus et al., 2018), a key enzyme controlling mitochondrial β-oxidation, was reduced in atrial myocardium in HFD compared to ND mice (Figure 2E) reflecting a higher CPT-1 activity in the former animal group. Taken together, these data unraveled a major metabolic remodeling in atrial myocardium in response to an obesogenic diet characterized by an accumulation of C18-containing lipids as well as long and polyunsaturated species, a shift to increased expression of genes involved energy metabolism pathways, such as glycolysis and β-oxidation, and an accumulation of energy substrates such as ketone bodies, glycolysis intermediates and long-chain TG.

### Distinct metabolic and lipidomic profiles between plasma and atrial myocardium in obese mice

There was a sharp separation of plasma metabolic profiles between HFD and ND mice as assayed by PCA analysis (Supplemental Figure S4A) with a marked decrease of individual lysophospholipids (LPE(16:0), (16:1), (20:5) and LPC(16:1), (18:1), (20:1), (20:5), and PI ((32:1), (34:1), (34:2), (36:4), (38:3)) species whereas individual PC ((18:0), (30:2), (38:4), (40:4)), PG ((36:2), (36:3), (38:4)), sphingolipid (SM ((32:1), (36:2), (42:3), (42:4)) and ceramides (Cer (d18:1/18:1), (18:1/20:0), (18:1/22:0), (18:2/22:0)) species were increased in the plasma of HFD mice (Supplemental Figure S4B). A marked decrease in carnitine (C0) and short-chain acylcarnitine (C2, C6) together with an elevation of long-chain acylcarnitine (C18) characterized the plasma metabolic phenotype of HFD mice (Supplemental Figure S4B). Enrichment in sphingolipids (DHC, ceramides and sphingomyelin), PC and PG, carbohydrates, amino acids, ketone bodies and polyols together with the depletion of PI and acylcarnitines of plasma from HFD mice were evidenced by an analysis of forest plots (Supplemental Figures S4C,D). These results indicate that the metabolic environment of HFD mice is mainly characterized by higher circulating amounts of sphingolipids and energy substrates such as carbohydrates, amino acids and ketone bodies.

Although there was a large overlap of lipidome between atrial myocardium and plasma, some alterations appeared specific to the tissue such as PS, PG, very short or very long TG species (Figure 3A). For instance, alterations of negatively charged PL (PI, PS and PG) were further exacerbated in the atrium. The increase in PE, PE and LPC, the decrease in very short TG and FFA and the increase in very long TG appeared as the specific lipidome of the atrial myocardium of HFD mice (Figure 2C and Figure 3C). On the other hand, despite difference in amplitude, there was an overlap of metabolite alterations between plasma and atrial myocardium (Figure 3B).

**Figure 3.**
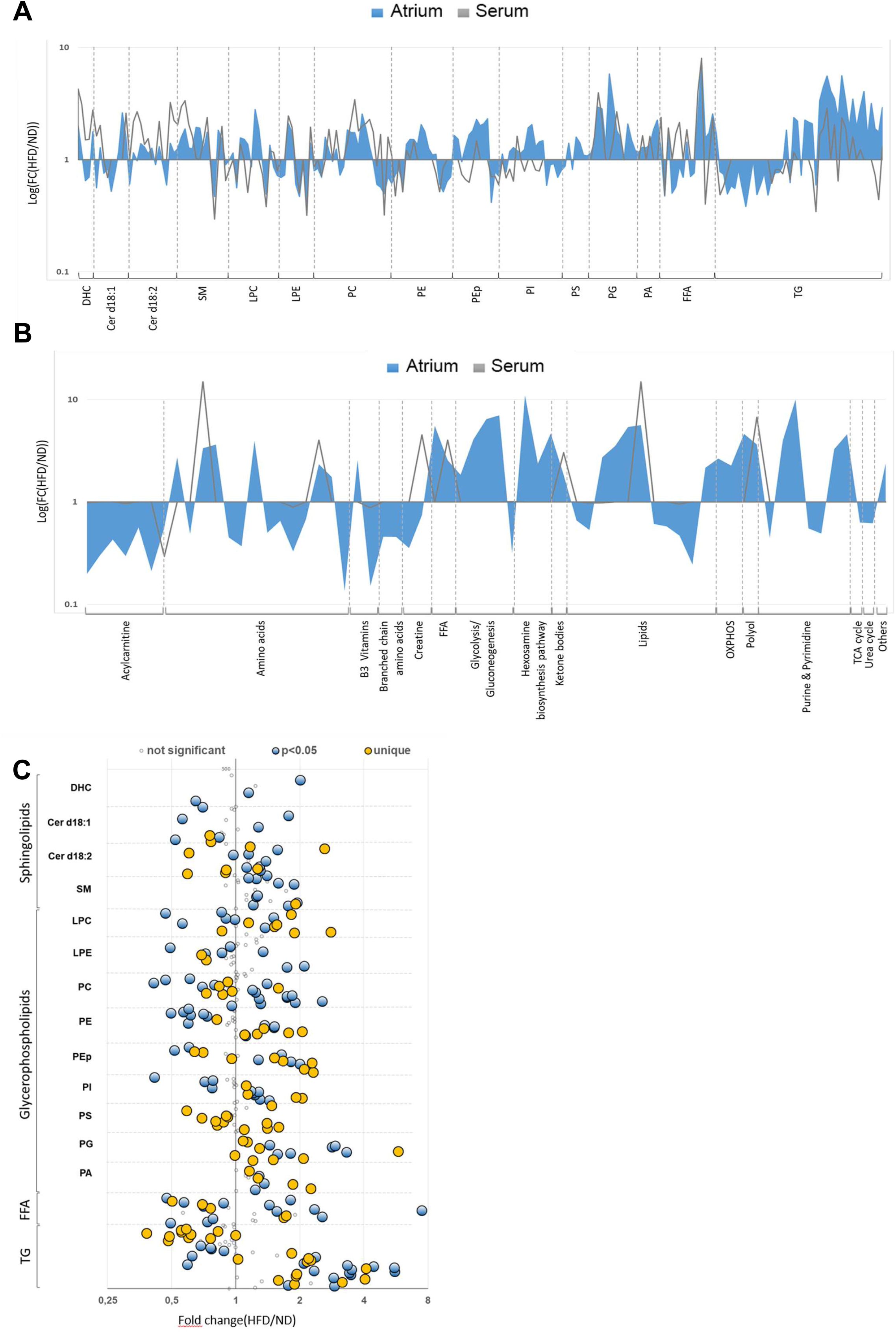
Distinct metabolic and lipidomic profiles between plasma and atrial myocardium after prolonged obesogenic diet. (**A**, **B**) Superimposition of lipidomics (**A**) and metabolomics (**B**) fold change plots of atrium (in blue) and serum (in grey) and showing large overlap between groups. P < 0.05, FDR corrected. (**C**) Forest plot showing individual lipids in atrial myocardium ordered by chain length and for each lipid subclass. Blue and yellow plots show significantly altered lipid species in HFD versus ND fed atria. In blue, plots common to atrium and serum, in yellow, plots specific to atrium. Data are expressed as mean ± SEM of independent experiments. Groups are compared using t-test, p value < 0.05 corrected by FDR.

Correlation analyses between individual lipid species in the atrium showed that most of ceramides correlated with short-chain TG containing SFA and MUFA, whereas an inverse correlation was observed between both these lipid classes and either TG with long fatty acid chains containing PUFAs (mostly AA) or PG (Supplemental Figure S5A-B and Figure S6). The identification of specific clusters of lipids that negatively or positively correlated pointed to the impact of circulating lipids on atrial myocardium lipidome (Supplemental Figure S7A). There was also a strong correlation between β-oxydation intermediates (carnitine and acylcarnitines) in atrium with those in plasma (Supplemental Figure S7B). Of note, glycolytic metabolites in the atrium were positively correlated to hexosamines and nucleotides, highlighting the use of carbons from the glucose molecule by hexosamine biosynthetic and pentose phosphate pathways at the expense of the glucose oxidation pathway.

### Measurement of mitochondria respiration revealed an increased β-oxidation in atrial trabeculae from obese mice

We subsequently examined whether this metabolic shift was associated with changes in respiratory properties. Mitochondrial respiration was assayed by measuring oxygen consumption in permeabilized atrial fibers. Mitochondrial respiration rates were not altered after the sequential addition of glutamate (complex I), pyruvate (production of acetyl-coA and NADH by PDH + complex I), succinate (complex I and II) and amytal to inhibit complex I (complex II) in both right and left atria of HFD mice eliminating major mitochondrial dysfunction (Figure 4A,B). Of note, after a TMPD-ascorbate addition, a modest but significant decrease in O_2_ consumption was observed only in the right atria of HFD mice (12.81±0.72 μmol O_2_/min/g dry weight *versus* 15.66±1.11 μmol O_2_/min/g dry weight in ND mice, p=0.045) (Figure 4A-B).

**Figure 4.**
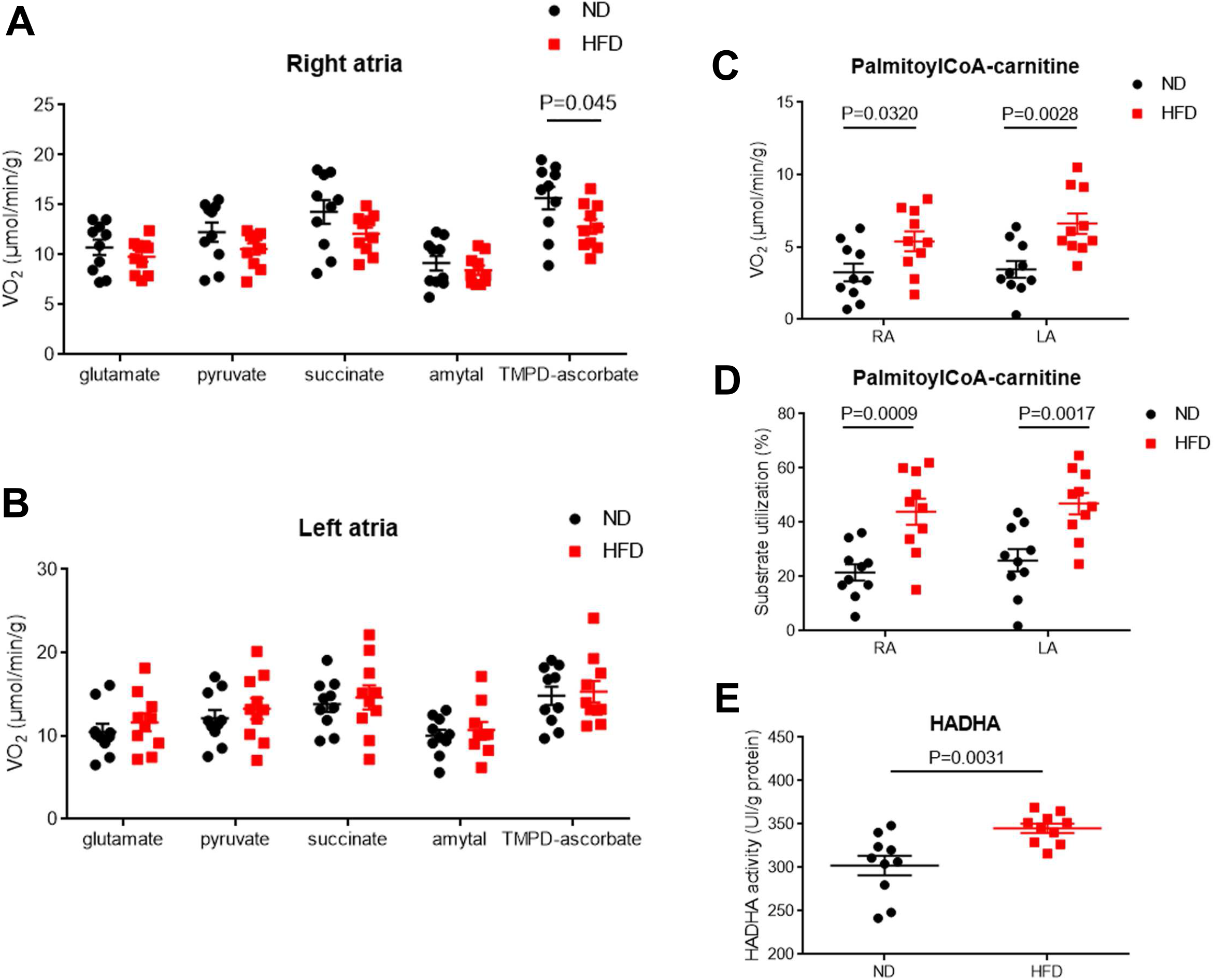
Measurement of mitochondria respiration revealed an increased β-oxidation in atrial trabeculae from obese mice. (**A, B**) Dot plots represent mitochondrial respiration rates in right (**A**) or left (**B**) atrial fibers permeable incubated with glutamate (10 mM), pyruvate (1 mM, complex-I substrate), succinate (15 mM, complex-II substrate), amytal (1 mM, complex I inhibitor) or TMPD-ascorbate (0.5 mM, complex IV activator) (n=10). (**C**) The O2 consumption was measured in both right and left atria of mice upon 16-we HFD after palmitoyl-CoA-carnitne (100 μM) addition (n=10). (**D**) The measure of substrate utilization after palmitoyl-CoA-carnitne (100 μM) addition was normalized with maximal O2 consumption (V_max_) in both left and right obesogenic mice atria (n=10). (**E**) The activity of hydroxyacyl-CoA deshydrogenase (HADAH) was measured in mice atria upon 16-we HFD or ND (n=10). Data are expressed as mean ± SEM of independent experiments. Statistical analysis was performed using unpaired t-test.

HFD induced an increase of O_2_ consumption after the addition of palmitoyl-CoA-carnitine (PCoA-Car) in both the left atrium (6.61±0.71 μmol O_2_/min/g dry weight *versus* 3.45±0.58 μmol O_2_/min/g dry weight in HFD and ND mice, respectively p=0.003) and the right atrium (5.39±0.69 μmol O_2_/min/g dry weight *versus* 3.25±0.61 μmol O_2_/min/g dry weight in HFD and ND mice, respectively, p=0.032) (Figure 4C). To evaluate substrate utilization, PCoA-Car was added and respiration rates were expressed as a percentage of maximal Complex I + II respiration rates. In HFD mice, a ~2-fold increase in oxygen consumption was observed in right atria (43.94±4.78 % *vs* 21.58±3 % in HFD and ND mice respectively, p=0.0009) and a 1.8-fold increase in left atria (46.9±3.96 % *versus* 25.99±4.11 % in HFD and ND mice respectively, p=0.0018) (Figure 4D), indicating that atrial myocardium mitochondria from HFD mice had a higher capacity to use fatty acids than that of ND mice.

Finally, the activity of the hydroxyacyl-CoA dehydrogenase (HADHA), a key enzyme of β-oxidation, was increased in the atria of HFD mice (345.14±5.40 IU/g *versus* 302.35±11.32 IU/g protein, HFD and ND mice respectively, p=0.0031) (Figure 4E). Taken together, these results indicate an activation of β-oxidation without major alteration of mitochondrial respiratory chain function in the atria of HFD mice.

### Action potential shortening and K-ATP current activation in the atrial myocardium of obese mice

As myocardial electrophysiology is tightly regulated by cellular metabolism, we compared the electrical properties of the atria between HFD and ND mice. First, propagated action potentials (APs) were recorded *ex vivo* in left atrial trabeculae, preparation using the gold standard glass-microelectrode technique. In HFD mice, AP duration (APD) was shortened at a different time of repolarization, for instance APD90 (69.46 ± 1.39 ms *vs* 76.98 ± 1.65 ms in HFD and ND mice, respectively (p<0.0001)) (Figure 5A). Of note, AP shortening was already observed after 2 weeks of HFD (Supplemental Figure S8).

**Figure 5.**
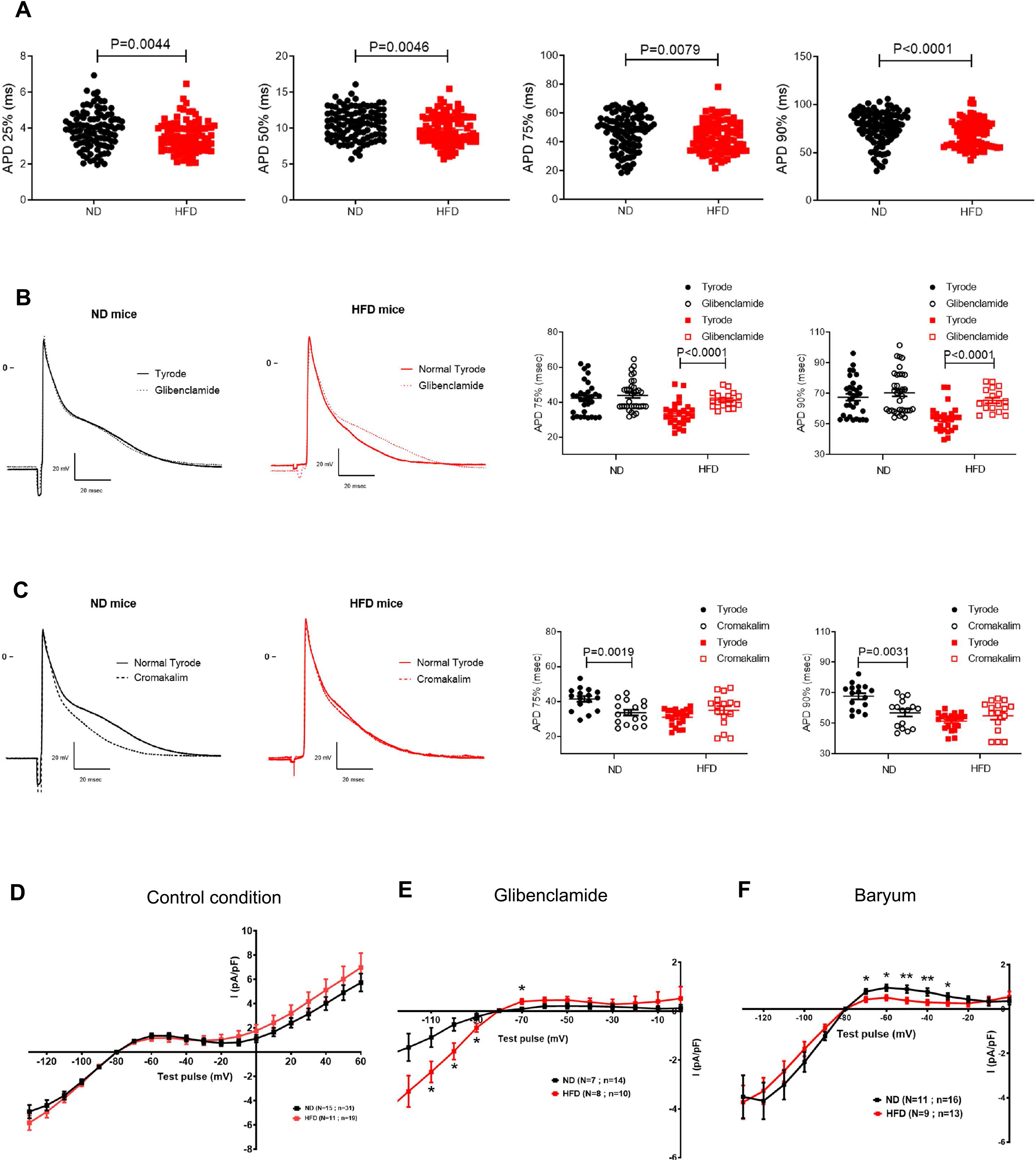
Action potential shortening and K-ATP current activation in the atrial myocardium of obese mice. (**A**) Action potential duration (ADP) measured at 25%, 50%, 75%, and 90% of the repolarization in left atrial trabeculae from HFD (red dot, n=9) and ND (black dot, n=11) mice. (**B,C**) Superimposition of action potential (AP) measured at 1Hz upon Tyrode and Glibenclamide (B) or Cromakalin (C) conditions and recorded after 16 week of diet in ND (black traces, n=10) and HFD (red traces n=10) mouse atrial trabeculae; right panels, quantification of the experiments by comparing the APD 75% and APD 90% between the various conditions tested. (**D**) Current-voltage relation of global potassium current measured at the end of 500 msec test pulse delivered form a holding potential of −80 mV and from −130 mV to +60 mV in perforated patch clamped atrial myocytes isolated form ND (black curve) and HFD (red curve) mice. (**E**) Using the same protocol as in panel D, current-voltage relation curves of subtracted component of potassium current suppressed by 5 μM Glibenclamide in ND (black curve) and HFD (red curve) mice. (**F**) Current-voltage relation curves of subtracted component of potassium current inhibited by 500 μM barium-sensitive current. Data are expressed as mean ± SEM of independent experiments. Statistical analysis was performed using Mann–Whitney U test (**A**) or 2-way ANOVA and Bonferroni post hoc test (**B-F**).

In order to examine the relationship between AP remodeling and the changes in myocardial metabolism described above, the effects of KATP modulators were tested on conducted AP in trabeculae from ND and HFD mice. Glibenclamide, a specific blocker of KATP channels, prolonged AP duration in HFD but not in ND mice, for instance at 75% (APD75: 41.28 ± 1.017 ms to 33.46 ± 1.365 ms, p<0.0001) and 90% of repolarization (APD90: 53.43 ± 1.669 ms to 65.41 ± 1.772 ms, p<0.0001) (Figure 5B). In contrast, cromakalim, a specific opener of ATP-sensitive potassium channels, shortened AP in ND but not in HFD mice (APD75: 41.49 ± 1.598 ms to 33.61 ± 1.653 ms, p =0.0034; APD90: 67.76 ± 2.041 ms to 56.83 ± 2.253 ms, p=0.0031) (Figure 5C).

These pharmacological profiles of APs strongly suggested an abnormal activation of KATP-dependent channels in atrial cardiomyocytes of HFD mice. This hypothesis was tested by recording potassium currents in isolated atrial myocyte by the patch-clamp technique. In wholecell configuration, no difference in potassium currents recorded between −130 mV to +60 mV was observed between HFD and ND atrial myocytes (Figure 5D). Note that in whole-cell configuration, the cytosolic medium is dialyzed by a pipette solution such that the cellular metabolic status is “normalized” during the current recording. Thus, in order to preserve the metabolic integrity of isolated atrial myocytes, potassium currents were recorded in the patch-perforated configuration (Akaike and Harata, 1994). The different components of the inward-rectifying potassium currents were then dissected using glibenclamide to block the *I*_K-ATP_ current and barium to block the *I*_K1_ background potassium current. As illustrated in Figure 5, the glibenclamide-sensitive component of the inward-rectifying potassium current was enhanced, whereas the barium-sensitive component was decreased in atrial myocytes isolated from HFD mice (Figure 5E,F). Taken together, these results indicate that K-ATP channels are activated in atrial myocytes, likely contributing to the AP shortening in HFD-fed mice.

### Acquisition of an adipogenic profile of the atria myocardium of obese mice

We next investigated the adipogenic profile of atrial tissue. We first looked for histological evidence of the presence of cardiac adipose tissue in HFD mice. Some adipose tissue depots were observed in atrio-ventricle grove and also in the epicardial area (Suffee et al., 2017) (Figure 6A). The adipose tissue score, defined as the proportion of atrial surface delineated with adipocyte (alveolar shape), indicated adipose tissue deposition had significantly increased in HFD mice compared to ND mice (Figure 6B). Furthermore, atrial myocardium of HFD mice showed a clear adipogenic phenotype as indicated by the increased expression at both protein and transcript levels of Perilipin-1 and fatty acid binding protein (Fabp) Fabp-4 (Figure 6C, D), and the increased mRNA levels for adiponectin (*Adipoq*), *Fabp-5* and cardiac specific *Fabp-3* (Figure 6E). Finally, the main adipogenesis transcription factors *C/ebpα* and *Pparγ* were also induced in atrial myocardium of HFD mice (Figure 6F).

**Figure 6.**
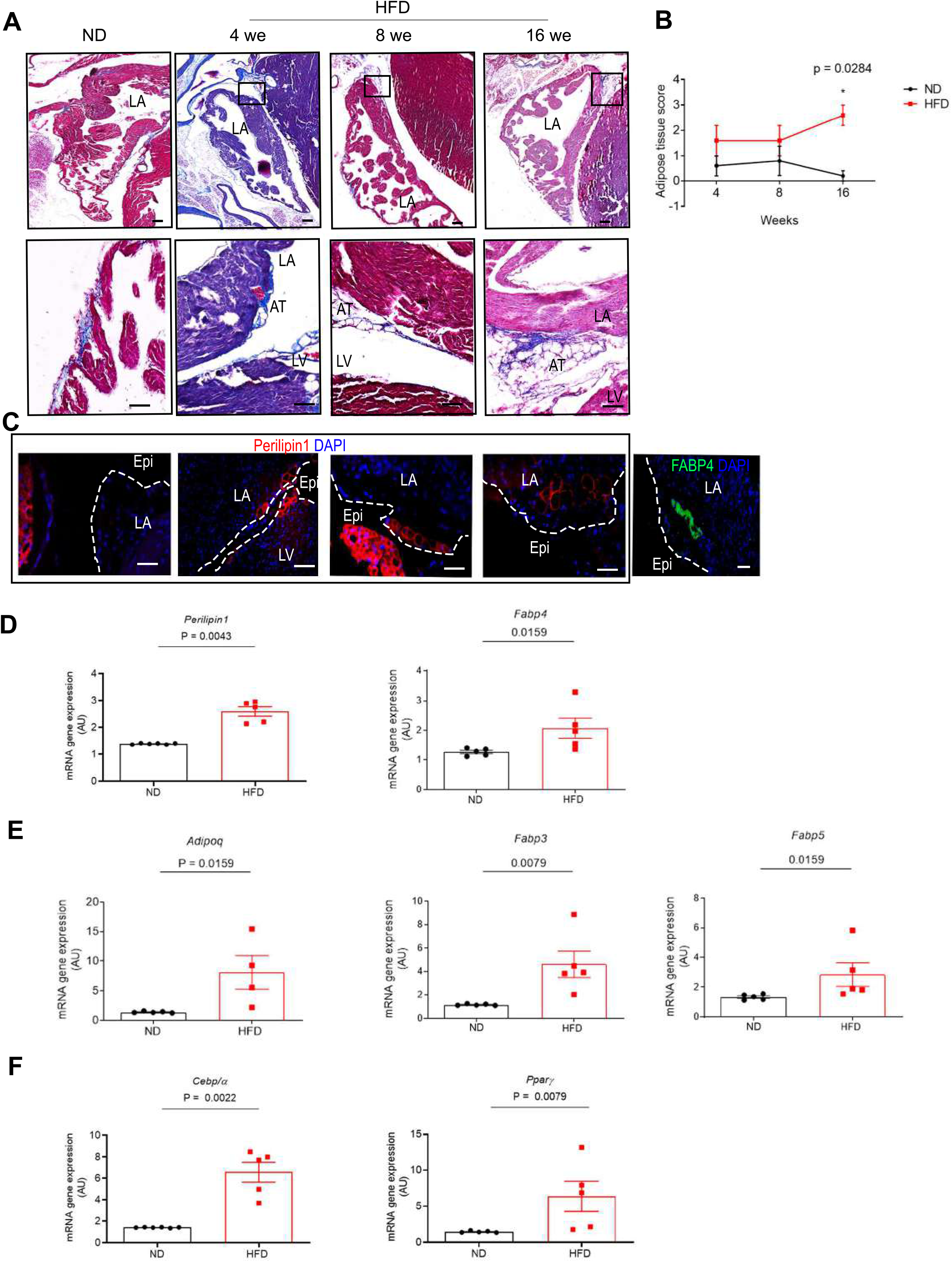
Acquisition of an adipogenic profile of the atria myocardium of obese mice. (**A**) Seven μm-thick sections of C57BL/6J mouse atrial tissue with 4we; 8we or 16we HFD or ND stained with Masson’s trichrome (n=5, each group). Scale bar, 100 μm, 50 μm. (**B**) Curves represent HFD or ND score of adipose tissue in C57BL/6J mouse atrial tissue upon 4we; 8we or 16we HFD or ND (n=5, each group). (**C**) Immunofluorescence for Perilipin (red) or FABP4 (green) in C57BL/6J mouse atrial tissue with 4we; 8we or 16we HFD or ND. Scale bar 50μm. (**D-F**) Transcript expression levels of adipogenic markers in C57BL/6J mouse atrial tissue upon 16-we HFD or ND (n=5). Data were normalized to *Gapdh* expression and are shown as fold change of treated cells as compared to untreated (UT) cells, and expressed in arbitrary units (AU). Data are expressed as mean ± SEM of independent experiments. Groups are compared with Mann–Whitney test.

Adipogenesis is characterized by a distinct inflammatory profile (Furuhashi et al., 2008) and also by the presence of chemokines in the secretome of atrial myocardium (Haemers et al., 2017; Suffee et al., 2017). Inflammatory cells could be detected in the atrial section of HFD mice (Figure 7A). Furthermore, tissue levels of mRNA of pro-inflammatory chemokines *Ccl2* and *Ccl5* were enhanced in HFD mouse atria (Figure 7B). Moreover, mRNA expression of genes encoding for proteins of immune cells *Cd45* in particular monocyte/macrophages *Cd14, f4/80, Cd68* and activated T-cells *Cd4, Pdcd1* (Figure 7C-D) were increased in the atria of HFD mice. Note that markers of T-cells *Cd8* and B-cells, *Cd19* were not induced in obesogenic mice atria (Figure 7C, D).

**Figure 7.**
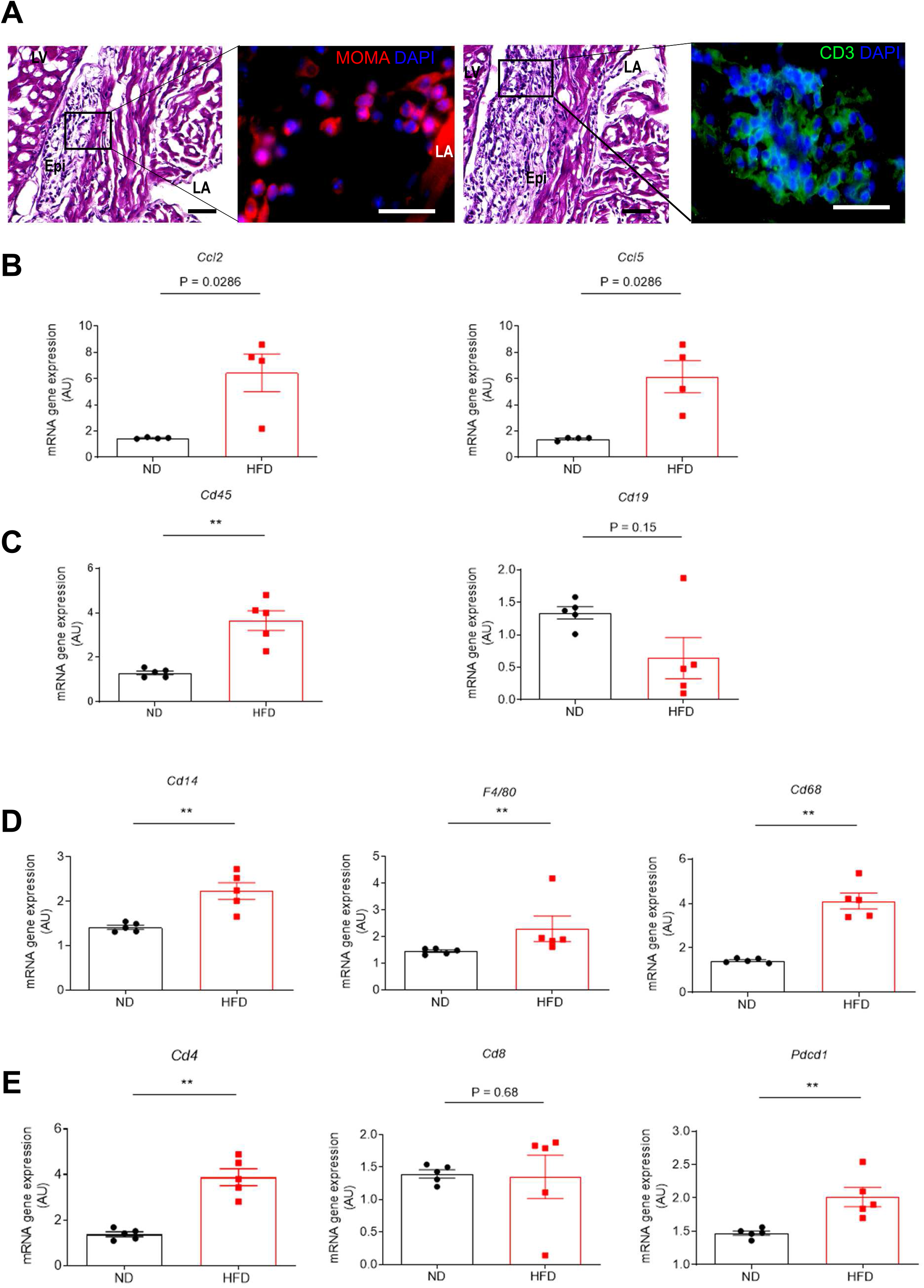
Evidence for an inflammatory profile of the atrial myocardium of obese mice. (**A**) Immunofluorescence staining of macrophages (MOMA) and T-cells (CD3) in freeze sections of atria from HFD (n=5). (Scale bars, 100μm, 50 μm). (**B-E**) Transcript expression levels of (**B**) pro-inflammatory chemokines gene *Ccl2* or *Ccl5*, (**C**) inflammatory cells genes *Cd45* and B-cells *Cd19*, (D) macrophages *F4/80*, *Cd68* and monocytes *Cd14*, or (**E**) T-cells *Cd8, Cd4* and *Pdcd1* in mice atria upon 16-weeks HFD (n=5). Data were normalized to *Gapdh* expression and are shown as fold change of treated cells as compared to untreated (UT) cells, and expressed in arbitrary units (AU). Data are expressed as mean ± SEM of independent experiments. Groups are compared with Mann–Whitney test, ** P = 0.0079. LA, left atria; LV, left ventricle; Epi, epicardium.

## DISCUSSION

Our study shows that an obesogenic diet causes profound transformation of the metabolic and lipidomic phenotypes of the atrial myocardium which acquires an adipogenic profile and changes its electrophysiology properties indicating the adaption of the atria to the new metabolic status. Diet can be a potent adipogenic factor for the atrial myocardium. Finally, this study provides new clues for understanding why metabolic disorders can be associated with an increased risk of AF.

Our study uncovers for the first time that during an obesogenic diet composed of a high amount of dietary fat, the atrial myocardium adopts an adaptive metabolic signature characterized by an impaired glycolysis, and an exacerbated auxiliary metabolic pathway generating hexosamines, nucleotides (through the pentose phosphate pathway) and polyols at the expense of the glucose oxidation and the use of BCAA. The increase of some glycolysis intermediates likely results from the accelerated gluconeogenesis in the atrium of HFD mice rather than increased glucose uptake through insulin-sensitive atrial glucose transporters, i.e. GLUT4 and GLUT8, which are impaired during diet-induced insulin resistance (Maria et al., 2018). In this vein, the content of palmitoleate (C16:1), a “lipokine” intended to preserve insulin signaling, (Cao et al., 2008) was drastically reduced in the atrial myocardium of obese mice. Such metabolic rewiring denotes a diabetic status of the atrium, classically observed in diabetic heart patients which was further supported with the increase of carbohydrates, ketone bodies and polyols.

Another major finding of our study is that an exacerbated flux of dietary lipids causes a drastic metabolic shift of the atrial myocardium toward other energy production pathways than glycolysis. For instance, metabolomics analysis and *ex vivo* testing of the mitochondrial respiratory chain indicate an increased FAO in atrial myocardium of HFD mice as illustrated by the reduction of the carnitine (C0)/acylcarnitine (C16) ratio, which reflects an increased CPT-1 activity and a more elevated capacity of atrial mitochondria to produce energy from palmitate. Both exacerbated, lipogenesis and FAO are also indicated by the upregulation of the activity of HADHA, a key enzyme of the β-oxidation pathway, and the up regulation of genes encoding for *Acaca* and *Fasn*. Interestingly, both myocardial FFA uptake and oxidation were demonstrated in obese, IR and diabetic patients using PET imaging with 1-^11^C-palmitate (Ussher et al., 2016). Lipidomics analysis of atrium appendages further support an increased FAO in the present study. A major remodeling of lipid metabolism was observed with enrichment in long and polyunsaturated lipids in most of the subclasses (TG, PC, PI, PS, PE and PE plasmalogens) at the expense of short and saturated lipid species of the corresponding subclasses. This was supported by an elevated C18 FA/C16 FA ratio in all lipid classes, highlighting an elongation mechanism. Interestingly, elovl6, an enzyme mediating the elongation of C16 FA to C18 FA and whose expression is induced by an HFD, was reported to promote lipogenesis, FAO and IR (Matsuzaka and Shimano, 2009). The enhanced elongation of C16 FA to C18 FA in the atrial myocardium might contribute to the increased FAO in HFD mice. This elongation mechanism also suggests a paralleled mechanism of lipid storage induction in response to the high flux of dietary fat further supported by the global enrichment in PG species, shown to trigger lipid storage in the adipose tissue in mice (Kayser et al., 2019a).

In addition to FAO, the atrial myocardium appears to use additional intracellular energy substrates such as ketone bodies as indicated by increase 3-hydroxybutyrate, the most abundant ketone body in atria of obese mice. Interestingly, proteomics and metabolomics analysis of human atrial appendages shows also a rise of β-hydroxybutyrate and ketogenic amino acids in AF patients (Mayr et al., 2008). Creatine levels are increased in the atria of obese mice. Creatine kinase protein levels, the key enzyme in the production of ATP from creatine, is also increased in atrial tissues from AF dogs (De Souza et al., 2010). Both ketone bodies and creatine could constitute an important fuel for the atrium to adapt to diet-driven metabolic stress or the fast beating rate of fibrillating as well as the hemodynamically overloaded ventricle (Ussher et al., 2016). Taken together, our data indicate that the atrial myocardium can activate compensatory metabolic pathways to overcome the reduction of energy production due to the defective glucose oxidation. In this way, the absence of an alteration of metabolites of the TCA cycle and the preserved mitochondrial respiratory activity are consistent with preserved oxidative metabolism and energy production of obese atria.

Further evidence of profound changes in the oxidative metabolism of the atrial myocardium of obese mice is the sustained and slight activation of K-ATP channels as evidenced by the short atrial AP with distinct sensitivity to K-ATP modulators and the increased IKATP in atrial myocytes in HFD mice. K-ATP channels are metabolic probe of the sub sarcolemma space, which are normally closed but open upon changes in the ATP/ADP ratio, metabolism alterations or changes in lipid content (Deutsch and Weiss, 1993). For instance, a preferential regulation of K-ATP channels by glycolysis upon lipolysis has been described in both cardiomyocytes and beta islets pancreatic cells and was attributed to close physical proximity of key glycolytic enzymes to the channels into the plasma membrane (Csonka et al., 2014; Deutsch and Weiss, 1993; Weiss and Lamp, 1989). A shift from glycolysis to predominant FAO as in this obesity model could reduce the availability of ATP for the K-ATP channels. The accumulation of long-chain CoA has been shown to prevent K-ATP channel closure and to directly activate channels (Liu et al., 2001; Schulze et al., 2003). Whatever the exact mechanisms underlying K-ATP channel activation, these findings indicate that energy substrate can regulate the electrical properties of atrial myocytes.

The other major finding of the present study is that atrial myocardium becomes adipogenic after a prolonged obesogenic diet as first indicated by the presence of adipose tissue at the epicardial surface of the atria or in the ventricle-atrial groove. Additionally, genes involved in adipogenesis such as *Perilipin and Adiponectin* (an adipokine secreted adipocyte) (Liao et al., 2005; Ouchi et al., 2011) or metabolism pathways such as transcription factors *C/ebpα, Pparγ* are upregulated in obese mouse atria. Finally, transcripts of lipid protein chaperones (FABP) including the adipocyte-type *Fabp4* (or *ap2*), the heart/muscle *Fabp3* and the epithelial/epidermal *Fabp5*, were detected in the atrial myocardium of obese mice (Lopez-Canoa et al., 2019; Shingu et al., 2018, 2020; Umbarawan et al., 2017). The FABPs facilitate the transport of FA to mitochondria (Rodríguez-Calvo et al., 2019) regulated lipid oxidation (Bonen et al., 2009) and storage (Rodríguez-Calvo et al., 2019). During myocardial hypertrophy, activation of the lipogenesis and increased fatty acid uptake are associated with an upregulation of *Fabp4* and *Fabp5* (Umbarawan et al., 2018). FABP proteins are also involved in the differentiation of adipogenic precursors derived from adipose tissue (Samulin et al., 2008). Both, FABP4 and FABP5 are increased at the early phase of adipogenesis and at the later phase for FABP3 (Samulin et al., 2008). A correlation between *Fabp* and *Pparγ* expression has also been reported during adipogenesis, pointing to a major adipogenic regulatory pathway (Fajas et al., 2002; Ren et al., 2002; Saladin et al., 1999).

The observation that obese mouse atria expressed transcripts of pro-inflammatory chemokines and markers of monocyte/macrophages and T-cells helper Cd4^+^ further supports the acquisition of an adipogenic phenotype of the myocardium. Indeed, cardiac adipose tissue is a major source of inflammatory mediators and contains a number of immune inflammatory cells (Iacobellis, 2015). For instance, T-cells CD4^+^ play a critical role in defending against obesity-associated metabolic disorders, notably by suppressing inflammation, thus maintaining the homeostasis of both immune responses and visceral adipose tissue (Zhu et al., 2017). Chemokines could be produced by both EAT (Haemers et al., 2017; Suffee et al., 2017) or pro-inflammatory AT macrophages (ATM-M1) (Mantovani et al., 2004). In obese mice, pro-inflammatory chemokines Ccl2 and Ccl5 could favor the recruitment of Cd14^+^ monocytes and the differentiation into mature macrophages expressing *F4/80* and *Cd68* within the atria. Of note, FABP4 is also induced during monocytes differentiation into macrophages in perivascular adipose tissue (Kazemi et al., 2005; Lamas Bervejillo et al., 2020). In line with those findings, an overall enrichment of PG species, which are inflammation-responsive lipids (Kayser et al., 2019b), was observed in atria of the HFD.

There are other circumstances during which atrial myocardium can acquire an adipogenic profile. For instance, the rapid rate of beating of sheep atria or the rapid pacing of isolated human atria induce the expression of an adipogenic gene profile (Chilukoti et al., 2015). We previously reported that the atrial natriuretic peptide secreted by atrial myocytes in response to stretch induces the differentiation of epicardial progenitors in adipocytes contributing to EAT expansion (Suffee et al., 2017). Collectively these studies suggest that the working conditions of the atrial myocardium can regulate the accumulation of EAT in line with the role of cardiac adipose tissue for fueling the myocardium with FA.

Several characteristics of the metabolically remodeled atrial myocardium can contribute to AF vulnerability of obese mice. For instance, EAT accumulation and inflammation are well established arrhythmogenic factors (Abed et al., 2013; Fontes-Carvalho et al., 2014; Kondo et al., 2018). Tissue accumulation of ketone bodies and sphingolipids observed in obese mice atria can be lipotoxic, contributing to myocardial dysfunction and activation of arrhythmogenic mechanisms (Goldberg et al., 2012) as reported during heart failure or in diabetic and obese patients (Sikder et al., 2018) (DHC, SM, Cerd 18:2). Whether these metabolic-driven electrical remodeling can contribute to AF vulnerability in patients, notably during metabolic disorders, remains to be elucidated.

## CONCLUSION

The present study indicates that a high fat diet induces profound and complex transformations of the metabolomic and lipidomic profiles of the atrial myocardium characterized by a predominance of FA oxidation and an increased lipid storage. This new metabolic phenotype is associated with acquisition of adipogenic traits of the atrial myocardium together with activation of metabolic-sensitive potassium channels indicating a new homeostasis of the atria. Further studies aiming to understand how alterations in the triangle linking metabolism, physiology and histology of the atrial myocardium contribute to the substrate of AF are now warranted.

## LIMITATIONS OF STUDY

Our results have been obtained in mice that can be submitted easily to drastic changes in diet and with a cardiac energy metabolism characterized by high glycolytic component, in sharp contrast with humans. However, there is evidence in humans as well of the impact of diet on energy metabolism and functional properties of the heart. For instance, in prediabetes patients, a postprandial increase in cardiac uptake of dietary fatty acid has been directly visualized using position emission tomography imaging and has been shown to correlate with impaired pump function and reduced myocardial energy efficiency (Labbé et al., 2011, 2012; Noll et al., 2015)

## ACKNOWLEDGMENTS

This work was supported by the French National Agency through the national program *Investissements d’Avenir* (Investments for the Future), grant ANR-10-IAHU-05 (to N.S, E.Bap G.D., F.I, M.P, ML, E.Bal and S.N.H.) and through the Project RHU-CARMMA, grant ANR-15-RHUS-0003 as well as the *Fondation de La Recherche Medicale* (Foundation of Medical Research) (to N.S. EBap, EBal and S.N.H.). This project received funding from the European Union’s Horizon 2020 Research Programme, grant 633193 “CATCH ME” (N.S. and S.N.H.).

## AUTHOR CONTRIBUTIONS

NS, EB designed, performed, analyzed and interpret most of the experiments and contribute to the writing of the manuscript. ML, FI designed performed, analyzed and interpreted the lipidomic and metabolomics experiments. ND, CP, VG designed, performed, analyzed and interpreted the electrophysiological experiments. GD performed, analyzed experiments. NM performed, analyzed and the clinical experiments in mouse model. LL contribute to the design, the interpretation and the writing of the manuscript. EB designed and interpreted the electrophysiological experiments and contributed to the writing of the manuscript. MM designed and interpret the respiration mitochondria experiments and contributed to the writing of the manuscript. WLG designed and interpret the metabolomics experiments and contributed to the writing of the manuscript. SNH designed the study, performed data interpretation and guidance of the project and write the manuscript.

